# Expanding the cultivable human archaeome: *Methanobrevibacter intestini* sp. nov. and strain *Methanobrevibacter smithii “*GRAZ-2*”* from human feces

**DOI:** 10.1101/2024.05.15.594450

**Authors:** Viktoria Weinberger, Rokhsareh Mohammadzadeh, Marcus Blohs, Kerstin Kalt, Alexander Mahnert, Sarah Moser, Marina Cecovini, Polona Mertelj, Tamara Zurabishvili, Jacqueline Wolf, Tejus Shinde, Tobias Madl, Hansjörg Habisch, Dagmar Kolb, Dominique Pernitsch, Kerstin Hingerl, William Metcalf, Christine Moissl-Eichinger

**Affiliations:** D&R Institute of Hygiene, Microbiology and Environmental Medicine, Medical University of Graz, Graz, Austria; BioTechMed Graz, Graz, Austria; Research Group Metabolomics, Leibniz Institute DSMZ-German Collection of Microorganisms and Cell Cultures GmbH, Braunschweig, Germany; Otto Loewi Research Center, Medicinal Chemistry, Research Unit Integrative Structural Biology, Medical University of Graz, Graz, Austria; Core Facility Ultrastructure Analysis, Medical University of Graz, Graz, Austria; Gottfried Schatz Research Center, Cell Biology, Histology and Embryology, Medical University of Graz, Graz, Austria; Department of Microbiology, University of Illinois, Urbana, Illinois, USA

**Keywords:** Methanobrevibacter smithii, *Methanobrevibacter intestini*, fecal methanogens, human archaeome

## Abstract

Two mesophilic, hydrogenotrophic methanogens, WWM1085 and *M. smithii* GRAZ-2 were isolated from human fecal samples. WWM1085 was isolated from an individual in the USA, and represents a novel species with in the genus *Methanobrevibacter*. *M. smithii* GRAZ-2 (= DSM 116045) was retrieved from fecal samples of a European, healthy female and represents a novel strain within this genus. Both *Methanobrevibacter* representatives form non-flagellated, short rods with variable morphologies and the capacity to form filaments. Both isolates showed the typical fluorescence of F_420_ and methane production.

Compared to *M. smithii* GRAZ-2, WWM1085 did not accumulate formate when grown on H_2_ and CO_2_. The optimal growth conditions were at 37°C, and pH 7. Full genome sequencing revealed a genomic difference of WWM1085 to the type strain of *M. smithii* PS (type strain; DSM 861), with 93.55% ANI and major differences in the sequence of its *mcrA* gene (3.3% difference in nucleotide sequence). Differences in the 16S rRNA gene were very minor and thus distinction based on this sequence might not be possible. *M. smithii* GRAZ-2 was identified as a novel strain within the *Methanobrevibacter* genus (ANI 99.04 % to *M. smithii* PS).

Due to the major differences of WWM1085 and *M. smithii* type strain PS in phenotypic, genomic and metabolic features, we propose *M. intestini* sp. nov. as a novel species with WWM1085 as the type strain (DSM 116060T = CECT 30992).

## Introduction

*Methanobrevibacter* species are widespread and have been found in numerous host microbiomes. They exhibit remarkable adaptability in engaging with both animal hosts and non-archaeal elements within their microbiome. By metabolizing diverse small fermentation byproducts, these species effectively facilitate and support various syntrophic interactions. They stand out as the predominant archaea thriving in the gastrointestinal tracts of not only several animals (1–3).

Among these species, *M. smithii* (with four isolates currently available, source: Global catalog of microorganisms, Nov 2023, https://gcm.wdcm.org/), represents the most prevalent archaeon within the human gut, exhibiting an average relative abundance of up to 2% in individuals with high methane emission levels in their breath (4).

It’s worth noting that the *M. smithii* type strain PS (DSM 861; hereby referred to as: “*M. smithii* PS”), was initially isolated from sewage samples rather than human feces (5). A contamination of the sewage sample with human feces cannot be excluded, in particular as the gastrointestinal tract is the most favorable habitat for *M. smithii*. In contrast, strain ALI (DSM 2375; hereby referred to as: “*M. smithii* ALI”) (6) is considered as one of the first publicly available *M. smithii* strains described and isolated directly from human fecal samples.

Additionally, a recent discovery indicated that *M. smithii* encompasses two distinct clades, tentatively labeled as smithii and smithii_A within the GTDB taxonomy (Rinke et al., 2021). This differentiation was further corroborated through genomic analyses and the incorporation of numerous metagenome-assembled genomes (MAGs) from studies on the human microbiome, confirming the taxonomic separation between smithii and smithii_A (8).

It was found that the median genome size of smithii_A slightly surpasses that of smithii (1.9 Mbp compared to 1.7 Mbp), while showing an average nucleotide identity (ANI) of 93.95%. Despite this variance, key genes linked to methanogenesis were shared between both strains. The *mcrA* gene exhibited an average amino acid sequence difference of 2.15% (8), a potential marker for distinguishing these clades using molecular methods (9). Following these observations, smithii_A was tentatively designated as a distinct species namely, *Candidatus* Methanobrevibacter intestini.

*Cand*. M. intestini is represented by WWM1085 (formerly recognized as a strain of *M. smithii*), which was initially isolated from human stool in the presence of CO_2_-H_2_ as a carbon and energy source (Chibani et al., 2022; Jennings et al., 2017). This species demonstrates extensive distribution and a notably high prevalence among the human population, accounting for approximately 90.01% ((11). In the present paper, we further describe *Methanobrevibacter intestini* as a novel species within the *Methanobrevibacter* genus using comparative 16S rRNA and genome sequencing, culture-based methods, electron microscopy, lipidomics, and metabolomics. We provide this *Methanobrevibacter intestini* strain (WWM1085) as a new addition to the culture collection (DSM 116060) of anaerobic archaea found in humans. Moreover, we characterize another newly isolated strain of *M. smithii* called *M. smithii* GRAZ-2.

## Materials and methods

### Sources of microorganisms

Strain WWM1085 (= DSM 116060 = CECT 30992) was enriched by the Department of Microbiology, University of Illinois, Urbana, Illinois, United States, from a fecal sample (Mayo Clinic Minnesota, biome number 101159) in the presence of CO_2_ and H_2_ as a carbon and energy source. Further details are provided in the draft genome sequence announcement by Jennings et al. (Jennings et al., 2017). The enrichment was subcultured and purified via antibiotic treatment to a pure culture in 2021 at the Medical University of Graz, Austria. In detail, the growth medium (MpT1, see below) was supplemented with streptomycin sulfate (10 mg/ml) and penicillin G potassium salt (10 mg/ml) at a volume ratio of 1:100 (0.2 ml of the antibiotics mixture in a volume of 20 ml of medium).

1. *M. smithii* GRAZ-2 was isolated from a stool sample of a healthy female aged 42 at Medical University of Graz, Graz, Austria in 2018 in presence of CO_2_ and H_2_ as a carbon and energy source. This strain is also currently available in DSMZ (= DSM 116045) (German Collection of Microorganisms and Cell Cultures GmbH, Braunschweig, Germany).
2. *M. smithii* ALI (= DSM 2375) was obtained from the DSMZ and was used for comparative analysis.

### Ethical approval

Sampling of the human fecal sample was evaluated and approved by the local ethics committee (27-151 ex 14/15). Before participation, the participant signed an informed consent.

### Enrichment and isolation of strain GRAZ-2

The stool sample was collected from a fresh fecal sample with an ESwab (COPAN Diagnostics Inc., Italy). The collection fluid, which keeps anaerobic microorganisms alive, was transferred to ATCC medium 1340 (MS medium for methanogens, see below), supplemented with ampicillin (100 µg/mL), streptomycin (100 µg/mL), tetracycline (10 µg/mL) and nystatin (20 µg/mL). Methane production in the culturés headspace was verified after visible growth (turbidity and microscopy) using a methane sensor (BCP-CH4 sensor, BlueSens).

Enrichment of methanogens was achieved via fluorescence-activated cell sorting (FACS) exploiting the auto-fluorescence of the cofactor F_420_. FACS was performed at the ZMF Core Facility Molecular Biology in Graz, Austria. For detection of the F_420_ fluorescence, the violet laser (405 nm) and the bandpass filter 450/40 of the FACSAria III system (Becton Dickinson) were used. During the short sorting process, cells were kept and sorted into reduced medium. 500,000 events were collected and re-grown in liquid MS medium (see below).

Subsequently, the culture was plated on solid MS medium (1.5 % agar, w/v) in Hungate tubes using the roll-tube method as described (12). A single colony was picked and re-grown in liquid media. To further ensure purity, serial dilutions were performed.

### Growth media

Standard archaeal medium (13) was used to grow all isolates, with some modifications. This medium contained the following constituents (l^-1^ distilled water): 0.45 g NaCl, 5 g NaHCO_3_, 0.1 g MgSO_4_.7H_2_O, 0.225 g KH_2_PO_4_, 0.3 g K_2_HPO_4_.3H_2_O, 0.225 g (NH_4_)_2_SO_4_, 0.060 g CaCl_2_.2H_2_O, 2 ml (NH_4_)_2_Ni(SO_4_)_2_ solution (0.1% w/v), 2 ml FeSO_4_.7 H_2_O solution (0.1% w/v in 0.1 M H_2_SO_4_), and 0.7 ml resazurin solution (0.1% w/v). These compositions were then supplemented with 1 ml of each 10x Wolfés vitamin and 10x mineral solutions (13). Media was then deoxygenated with N_2_ and 0.75 g L-cysteine was added under anaerobic conditions. pH was adjusted to 7.0 if necessary. 20 ml of liquid was then aliquoted into 100 ml serum bottles, sealed with rubber stopper and aluminum cap, and pressurized with H_2_/CO_2_ (4:1) before autoclaving. Before use, 0.001 g/ml of yeast extract and 0.001 g/ml sodium acetate were added to the media.

For growth of WWM1085, MpT1 medium, based on AM-5 (14), was used with some modifications. Modified MpT1 medium had the following compositions (l^-1^ distilled water): 1 g NaCl, 0.5 g KCl, 0.19 g MgCl_2,_ 0.1 g CaCl_2_ x 2 H_2_O, 0.3 g NH_4_Cl, 0.2 g KH_2_PO_4,_ 0.15 g Na_2_SO_4_, 2g casamino acids, 2 g yeast extract, 0.082 g sodium acetate. Then, 1 ml trace element solution (1.2 ml HCl (12.5 M), 0.01 g MnCl_2_ x 4 H_2_O, 0.019 g CoCl_2_ x 6 H_2_O, 0.0144 g ZnSO_4_ x 7 H_2_O, 0.0002 g CuCl_2_ x 2H_2_O, 0.003 g H_3_BO_3_, 0.0024 g NiCl_2_ x 6 H_2_O, 0.0036 g Na_2_MoO_4_ x 2 H_2_O in 150 ml distilled water), 20 μl of selenite-tungstate solution (2 g NaOH, 0.01 g Na_2_SeO_3_.5H_2_O, and 0.017 g Na_2_Wo_4_.2H_2_O dissolved in 50 ml distilled water) and 0.7 ml resazurin solution (0.1% w/v) were added. Media was deoxygenated with N_2_ and subsequently, 0.24 g L-Cysteine and 2.52 g NaHCO_3_ were added. 20 ml of medium was distributed in 100-ml serum bottle and was then sealed and pressurized with H_2_/CO_2_ (4:1). After autoclaving, 0.2 ml of the following were added to each bottle under anoxic conditions: methanol (50 mM), dithiothreitol (0.154 g/L), Na-formate (0.034 g/L) / Na-coenzyme M (0.01 g/L) and vitamin solution (3 mg Biotin, 3 mg Folic acid, 15 mg Vitamin B6, 7.5 mg Vitamin B1, 7.5 mg Vitamin B2, 7.5 mg Nicotinic acid, 7.5 mg DL-Panthothenic acid, 1.5 mg Vitamin B12, 7.5 mg P-Aminobenzoic acid, 0.3 g Choline chloride dissolved in 150 ml distilled water) were added to each bottle under anaerobic conditions. The pH was adjusted to 7.0 if applicable. All growth experiments were carried out in triplicates under static conditions at 37 °C unless mentioned otherwise.

### Scanning electron microscopy (SEM)

For scanning electron microscopy, cells were mounted on coverslips, fixed with 2 % (w/v) paraformaldehyde in 0.1 M phosphate buffered saline, pH 7.4 and 2.5% glutaraldehyde in 0.1 M phosphate buffered saline, pH 7.4, and dehydrated stepwise in a graded ethanol series. Samples were post-fixed with 1 % osmium tetroxide for 1 h at room temperature and subsequently dehydrated in graded ethanol series (30-96 % and 100 % (v/v) EtOH). Further, Hexamethyldisilane (HMDS (Merck | Sigma - Aldrich) was applied. Coverslips were placed on stubs covered with a conductive double coated carbon tape. The images were taken with a Sigma 500VP FE-SEM with a SEM Detector (Zeiss Oberkochen) operated at an acceleration voltage of 5 kV.

### Optimum pH and temperature

The 100-ml serum bottles, each containing 20 ml of modified standard archaeal medium (for all tested isolates) and MpT1 (only for WWM1085), were inoculated with 2.5% (v/v) fresh cultures. Growth, measured in terms of OD at 600 nm, and methane production were monitored daily for 10 days to assess the impact of pH and temperature on growth. Methane levels were quantified using a gas sensor (BluSens, Germany), and data integration and analysis were performed using the provided BacVis Gas Formation software.

To explore the effect of pH on growth, media with different pH values ranging from 5 to 8 with 0.5 intervals, as well as pH values of 9, 10, and 11, were prepared by adjusting with varying amounts of 0.1 M NaOH or 0.1 M HCl. The pH values of the media were checked daily for potential alterations (pH indicator strips, VWR, Germany) and were maintained constant. The optimum pH was determined at 37°C.

For the determination of the optimum temperature, cultures were incubated at various temperatures (20, 30, 35, 37, 39, 40, and 50 °C), while pH was kept constant at 7. Temperature was monitored continuously using a temperature logger (Sensor Blue, Brifit) inside the incubator. Both pH and temperature experiments for WWM1085 were conducted in modified standard archaeal medium and MpT1.

### Culture purity check and sequencing

Cultures were routinely checked for purity using microscopy, PCR and Sanger sequencing. Microscopic examination of the cells was performed using a Nikon microscope equipped with a fluorescence attachment and a UV excitation filter. Extracted DNA was subjected to PCR targeting the archaea *mcrA* (forward primer sequence: 5’ CAACCCAGACATTGGTACTCCT 3’, reverse: 5’ GCTGGGGTGATGACAGTTCT 3’) and the bacterial 16S rRNA gene (primers 341F and 1391R) (15,16). Media blanks and no-template controls served as negative controls.

### Nanopore sequencing

The studied archaeal species underwent Nanopore sequencing using the MinION Mk1C system (Oxford Nanopore Technologies plc., UK) according to the protocols as detailed in (nanoporetech.com). To summarize, DNA extraction was done according to the manufacturer’s protocol (Invitrogen^TM^ PureLink^TM^ Microbiome DNA Purification Kit, Thermo Fisher Scientific Inc, USA), and subsequently, Nanodrop 2000c spectrophotometer (Thermo Fisher Scientific Inc., USA) and an Invitrogen^TM^ Qubit^TM^ 3 Fluorometer (Thermo Fisher Scientific Inc., USA) were used to confirm the quality and concentration of the extracted DNA. In addition, gel electrophoresis was employed for checking DNA fragmentation. DNA was then stored at −20 °C for further analyses.

In the process of preparing the library, DNA underwent repair utilizing the NEBNext Companion Module (New England Biolabs GmbH, GER). Subsequently, it was prepared for sequencing on a chemistry version 14 flow cell (R10.4.1, FLO-MIN114) following the Ligation sequencing gDNA – Native Barcoding Kit 24 V14 (SQK-NBD114.24) as outlined by the guidelines of the manufacturer as detailed in (nanoporetech.com).

### DNA-based comparisons

16S rRNA genes were retrieved from isolates’ genomes using the ContEst16S tool created by EzBioCloud (17) (Supplementary Table S1). The genomes were retrieved from own sequencing (GRAZ-2) or public databases (NQLD00000000 for WWM1085, all other accession numbers are provided in Supplementary Table S1).

Some genomes contained multiple copies of the 16S rRNA genes; in such a case, all were included in the subsequent analyses. Alignment of the sequences was performed via Muscle ((18,19), implemented in Mega11 (standard settings of MEGA11; (20)). All aligned 16S rRNA genes were manually trimmed to the same length. Pairwise distance estimation was performed using the standard settings. The matrix is available in Supplementary Table 2. The 16S rRNA gene-based tree was created via SILVA SINA using the FastTree option (model: GTR, rate model for likelihoods: Gamma; variability profile: Archaea; Positional variability filter, domain Archaea) (21–23).

*mcrA* genes were retrieved through MAGE genoscope ((24); Supplementary Table 3), a platform for genomic comparison. Genes were aligned through Muscle (see above) and pairwise distance estimation was calculated using the standard settings of Mega11. The matrix is provided in Supplementary Table 4.

The probability of whether one or two isolated *Methanobrevibacter* strains represent novel species was tested using JSpeciesWS (25). The Average Nucleotide Identity (ANI) was calculated against all isolates listed in Supplementary Table 5-6, and those provided by the curated reference database GenomesDB. This tool also provided the GC content of each genome.

The whole genome tree was calculated using MAGE genoscope (24) and the integrated “Clustering Genomes” function. The tree is constructed from the Mash distance matrix (26,27) and computed dynamically using a rapid neighbour joining algorithm. For details, please refer to the tutorial of MAGE genoscope.

### Lipid and carbohydrate profile analyses by mass spectrometry

Intact polar lipids were extracted from freeze-dried material (approx. 30 mg) using a modified Bligh and Dyer extraction as described previously (28–31). Briefly, two extractions each were performed using methanol/dichloromethane (DCM)/50 mM phosphate buffer pH 7-8 (2:1:0.8 v/v/v) and methanol/DCM/0.3 M trichloroacetic acid pH 2-3 (2:1:0.8 v/v/v). Combined supernatants were adjusted to a ratio of methanol/DCM/50 mM phosphate buffer of (2:1:0.9 v/v/v) by adding DCM and phosphate buffer, before the DCM phase was collected. The remaining mixture was additionally extracted twice with DCM and the combined DCM phases were evaporated to dryness. For HPLC-MS/MS analysis dried extracts were recovered in hexane/isopropanol/water (718:271:10 v/v/v).

Archaeal lipids were separated on a YMC-Triart Diol column (150 x 2.0 mm, 1.9 µm particles) and analyzed in positive ESI mode by mass spectrometry on an Agilent 6545 Q-ToF mass spectrometer (Agilent, Waldbronn, Germany) as described previously (28,32). Mass spectra were recorded in the mass range of m/z 300-2000. Core lipids were identified by the exact masses of their [M+H]^+^ ions.

For analysis of lipid associated sugars, lipid extracts were prepared as described above and hydrolyzed according to (33) with slight modifications. Briefly, dried extracts were resolved in 1 ml 2 N H_2_SO_4_ and incubated for 2 h at 100 °C. Afterwards the samples were chilled on ice and neutralized by adding 2 N NaOH (final pH 6-8). After centrifugation the supernatant was evaporated to dryness.

For GC-MS analysis of sugar residues tried extracts were reconstituted in 1 ml methanol and filtered through a Nylon spin filter to remove excess salt. The remaining supernatant was mixed with 10 µl of a 4 % ribitol-methanol solution and dried under a stream of nitrogen. In addition, non-hydrolyzed extracts were analyzed to detect any residual free sugars in the lipid extracts. Derivatization and GC-MS analysis was performed as described previously (34). Data analysis was performed with the MetaboliteDetector software (35) as described previously (36).

### Quantification of metabolic activity by NMR spectroscopy

Five replicates for each of the studied archaeal cultures (in MS medium and yeast extract supplement (see above)) at different time points (72h, 168h, and 240h), were subjected to analysis utilizing Nuclear Magnetic Resonance (NMR) spectroscopy, following the methodology outlined before (4). Briefly, a methanol-water mixture (2:1) was employed to eliminate proteins from samples followed by centrifugation. Subsequently, the supernatant was lyophilized, re-dissolved in sodium phosphate-buffered NMR buffer also containing 4.6 mM 3-trimethylsilyl propionic acid-2,2,3,3,-d4 sodium salt (TMSP) as internal standard, and subsequently transferred to NMR tubes. NMR analysis was then conducted on a Bruker Avance Neo NMR spectrometer running at 600 MHz and equipped with a TXI probe head at 310 K and Topsin 4.3 software (Bruker GmbH, Rheinstetten, Germany). The obtained spectra (cmpgpr1d/Carr-Purcell-Meiboom-Gill pulse sequence with 128 scans) were further processed using MATLAB 2014b (Mathworks, Natick, MA, USA), aligned, and normalized by probabilistic quotient normalization (37,38). For absolute quantification of carbonic acids, known peaks of substances of aligned raw spectra were integrated using trapezium subtraction for baseline correction (39), and eventually normalized on their respective proton number, J-coupling pattern, and TMSP integral of the sample in order to calculate their molar concentrations.

## Results and discussion

Based on our findings, the investigated archaeal strains, namely WWM1085 and *M. smithii* GRAZ-2, along with *M. smithii* DSM 2375, which was used for comparison, exhibit unique features discerned through our culture-based, genomics, and metabolomics methods. These distinctive characteristics are outlined in Supplementary Table 1 and elaborated upon in the subsequent sections.

### Morphology

WWM1085 cells appeared morphologically similar to both *M. smithii* ALI and *M. smithii* GRAZ-2 albeit being slightly shorter. In general, they measure 0.16-0.43 µm in width and 0.29-0.54 µm in length and appear mostly in the form of short rods with rounded ends (Fig. 1). Similar to *M. smithii* ALI and *M. smithii* GRAZ-2, not only they occurred in single cells, but were also observed more frequently in pairs, short chains or long filaments. Pili or flagella were not detected, but some cells appeared fluffy on their surface. All isolates showed F_420_ fluorescence, which is typical for methanogenic archaea, when observed under fluorescence microscopy (excitation 420 nm). No cells were observed in media controls.

**Fig 1.**
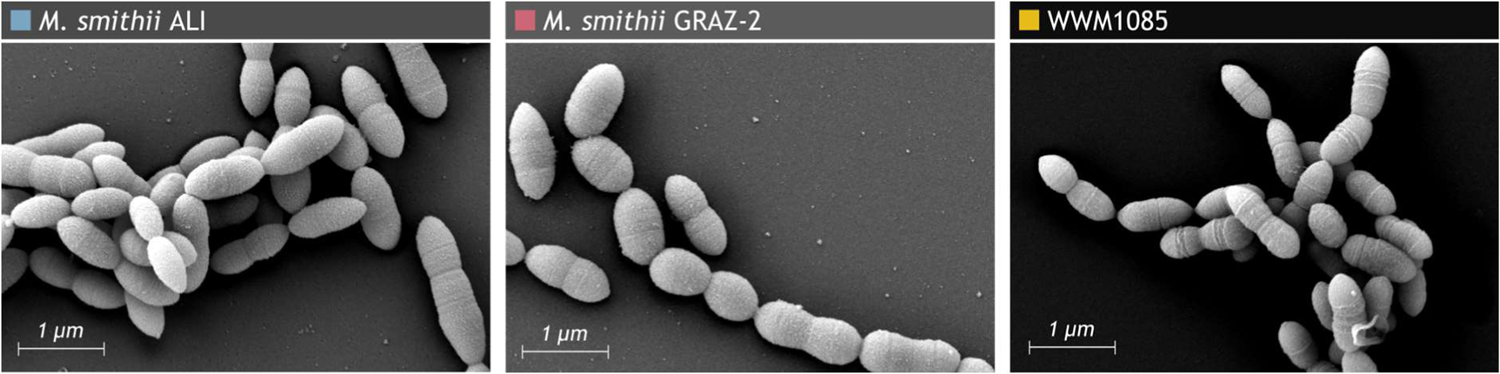
Scanning electron micrograph of *Methanobrevibacter smithii* ALI, *Methanobrevibacter smithii* GRAZ-2, and WWM1085.

### Substrates and nutritional requirements

WWM1085 underwent growth testing in two media, namely MS and MpT1 media, with various substrates to assess potential variations in its nutritional requirements compared to *M. smithii* ALI and *M. smithii* GRAZ-2. At pH 7 and 37°C, this strain demonstrated optimal growth in both media and reached high cell density after 72 hours (2.5% (v/v) inoculation), utilizing H_2_/CO_2_ as its energy source. No growth was observed when growth media were exposed to oxygen.

### Optimum pH range for growth and methane production

WWM1085 (in both media), along with *M. smithii* ALI, and *M. smithii* GRAZ-2 constantly produced methane across a broad pH spectrum (6.5 - 10) (Table. 1). On the basis of methane production and OD600, the optimum pH was found to be 7-7.5 in the MS medium. In modified MpT1, WWM1085 showed the optimal growth at a pH range of 6.5-7. The type strain *M. smithii* PS showed an optimal pH range between pH 6.9 and 7.4 (40). None of the isolates, showed growth at pH 5 and 11.

### Optimum temperature range for growth and methane production

All three isolates exhibited growth and methane production within a temperature range of 30-40°C (30, 35, 37, 39, and 40°C) (Table 1). In the modified standard archaeal medium, the optimum temperature range for *M. smithii* ALI, *M. smithii* GRAZ-2, and WWM1085 was determined to be 39-40°C, 39-40°C, and 35-39°C, respectively (Table 1). Notably, WWM1085 demonstrated the identical optimal growth at 35-39°C in the modified MpT1 medium, too. In summary, WWM1085 displayed a broader but lower temperature range for moderate or optimal growth as compared to the other two strains. No growth was observed under more extreme temperature conditions (20°C or 50°C) for any of the isolates. The type strain *M. smithii* PS showed an optimal temperature range between 37 and 39°C (40).

**Table 1.**
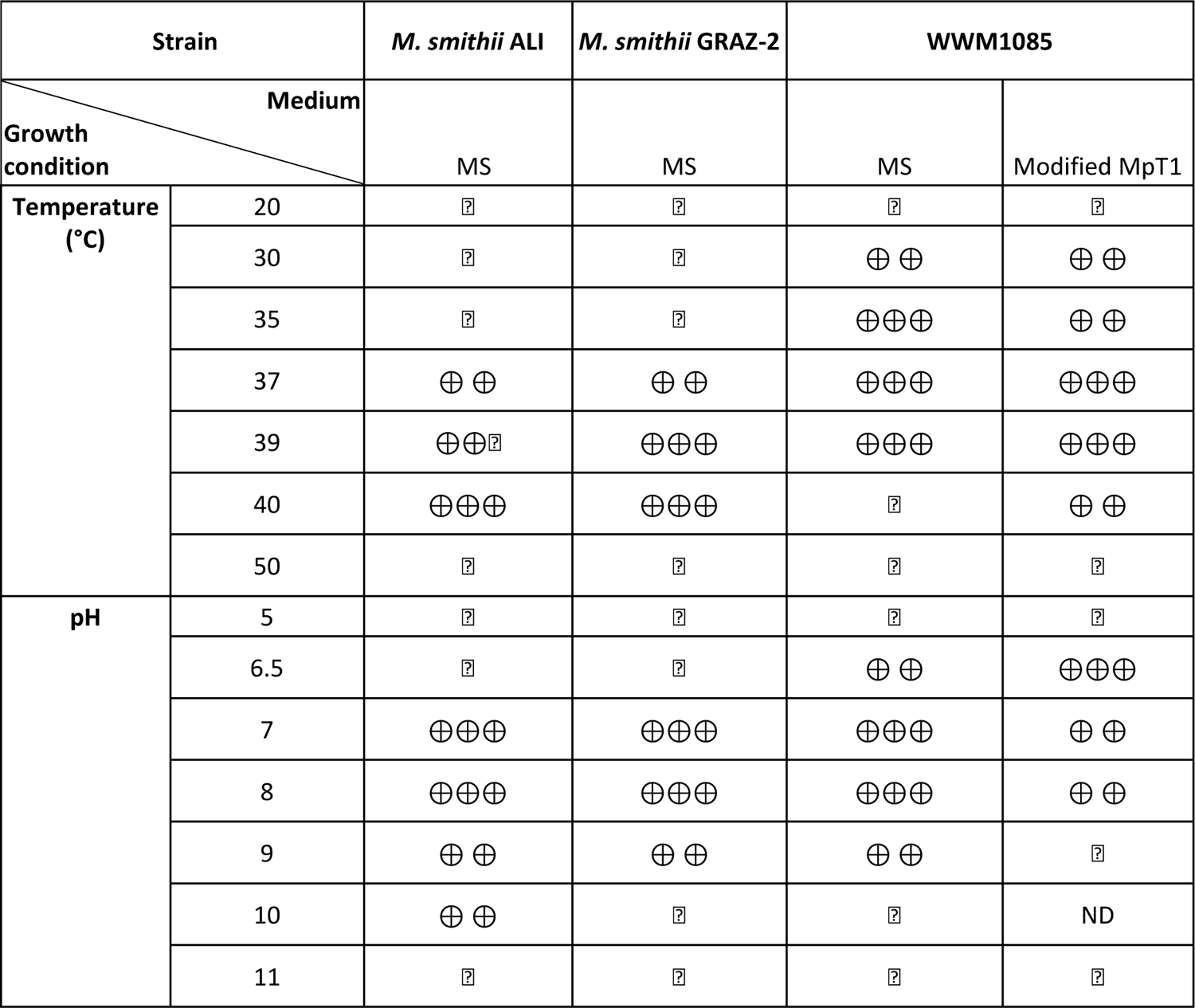
Optimal pH and temperature for the growth of the studied strains. Growth was determined by measuring OD600 in the growth medium and the ability of the strains to produce methane. (+) minimal growth; (++) moderate growth; (+++) optimal growth; (-) no growth; ND: Not determined.

### Phylogenetic relationships

The full-length 16S rRNA gene analysis of WWM1085 showed small variations as compared to closely related *Methanobrevibacter smithii* isolates (*M. smithii* ALI: 0.135%, *M. smithii* PS: 0.203%; Supplementary Table S2). These slight discrepancies pose a challenge for differentiating the isolates solely through 16S rRNA gene sequencing. It’s important to highlight that these sequence variations arose within a homopolymeric sequence region (multiple T), and at this juncture, we cannot dismiss the possibility of differences arising from sequencing artifacts or technical issues. The 16S rRNA gene of *M. smithii* GRAZ-2 was found to be highly similar to the genes of M. *smithii* PS and *M. smithii* ALI (difference: 0.068%; Supplementary Table S2) (Fig. 2).

**Fig. 2.**
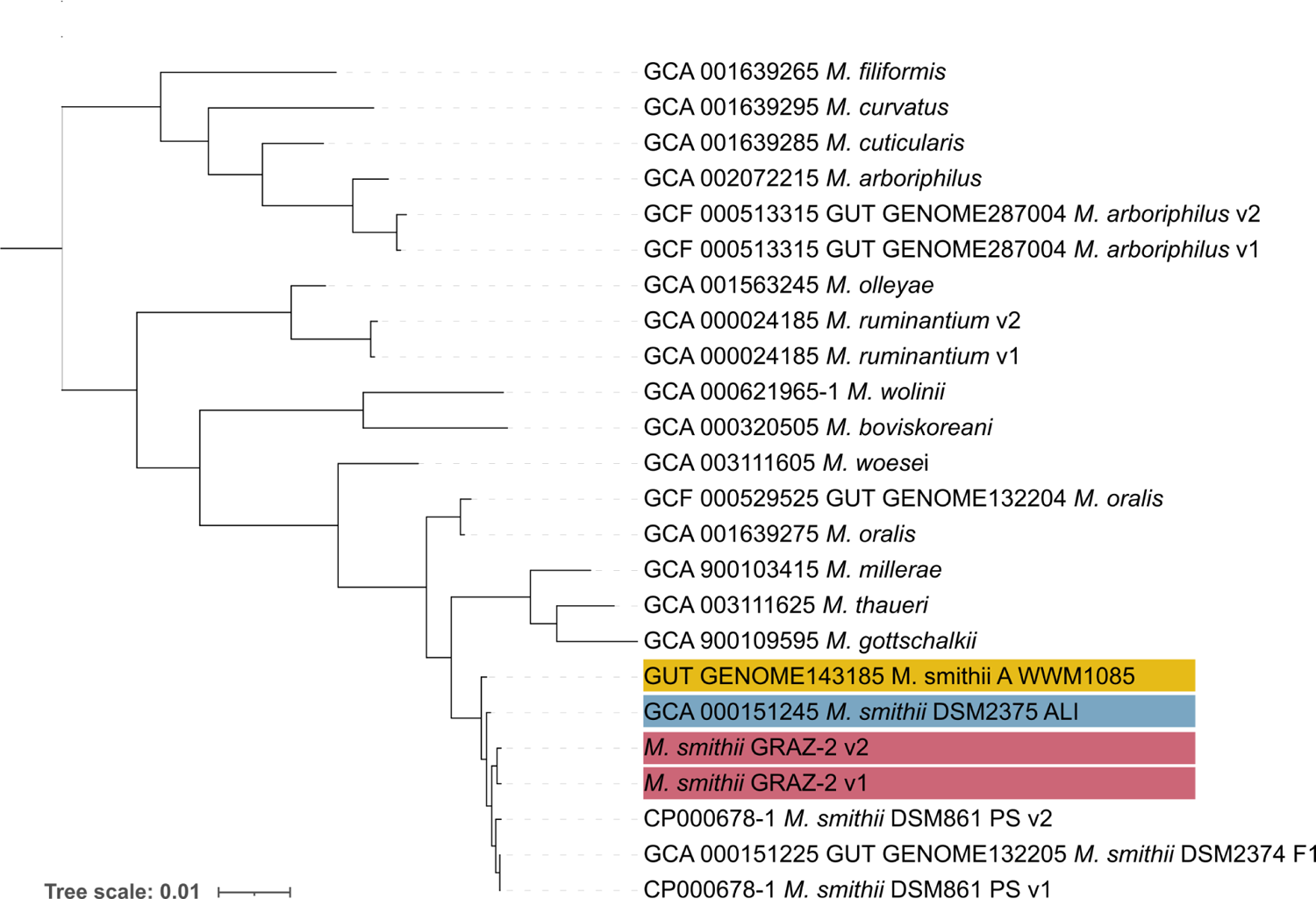
Phylogenetic relationship of the *Methanobrevibacter* isolates (WWM1085 in yellow, *M. smithii* ALI in blue, and *M. smithii* GRAZ-2 in pink) based on 16S rRNA gene analysis. *Methanosphaera stadtmanae* was used as an outgroup. Taxa fields contain the accession number (GenBank, UHGG (41), DSM). V1-4 refers to different versions of the 16S rRNA gene within the same genome. The tree was calculated via SILVA SINA using the FastTree option (see Materials and Methods).

Differences were more pronounced at *mcrA* gene level. At the nucleotide level, a minimum of 3% difference was observed between WWM1085 and *M. smithii* DSM 2374 F1, *M. smithii* ALI and *M. smithii* PS. On the other hand, *M. smithii* GRAZ-2 showed a difference of 0.0604 to 0.242% to the aforementioned strains, and therefore showed a higher similarity (Supplementary Tables S3-4). On amino acid level, the mcrA gene of WWM1085 showed a difference of 0.0703% to *M. smithii* DSM 2374.

Instead of employing DNA-DNA hybridization, comparisons based on full genome ANI were conducted (all genome-associated information is given in Supplementary Table S5). Notably, WWM1085 exhibited a slightly lower GC content as compared to *M. smithii* strains from the DSMZ collection (30.3% as opposed to 31.0-31.3%). When performing pairwise ANI calculations, the similarity values of WWM1085 against strains from GenomesDB and the culture collection consistently fell well below the species threshold (cutoff: 95%). The closest relatives were found to be *M. smithii* ALI (ANI: 93.04%) and *M. smithii* PS (ANI: 93.55%).

Consequently, it can be concluded that WWM1085 represents a distinct species within the *Methanobrevibacter* genus. However, it is important to note that despite these genomic differences, the disparities in the 16S rRNA gene are subtle, and in some cases, imperceptible in amplicon-based studies. *M. smithii* GRAZ-2 showed the closest relationship to *M. smithii* PS (ANI: 99.04 %) and therefore does not represent a novel species within the *Methanobrevibacter* genus (all information given in Supplementary Table S6). For visualization, a genome-based tree is provided in Fig. 3.

**Fig. 3.**
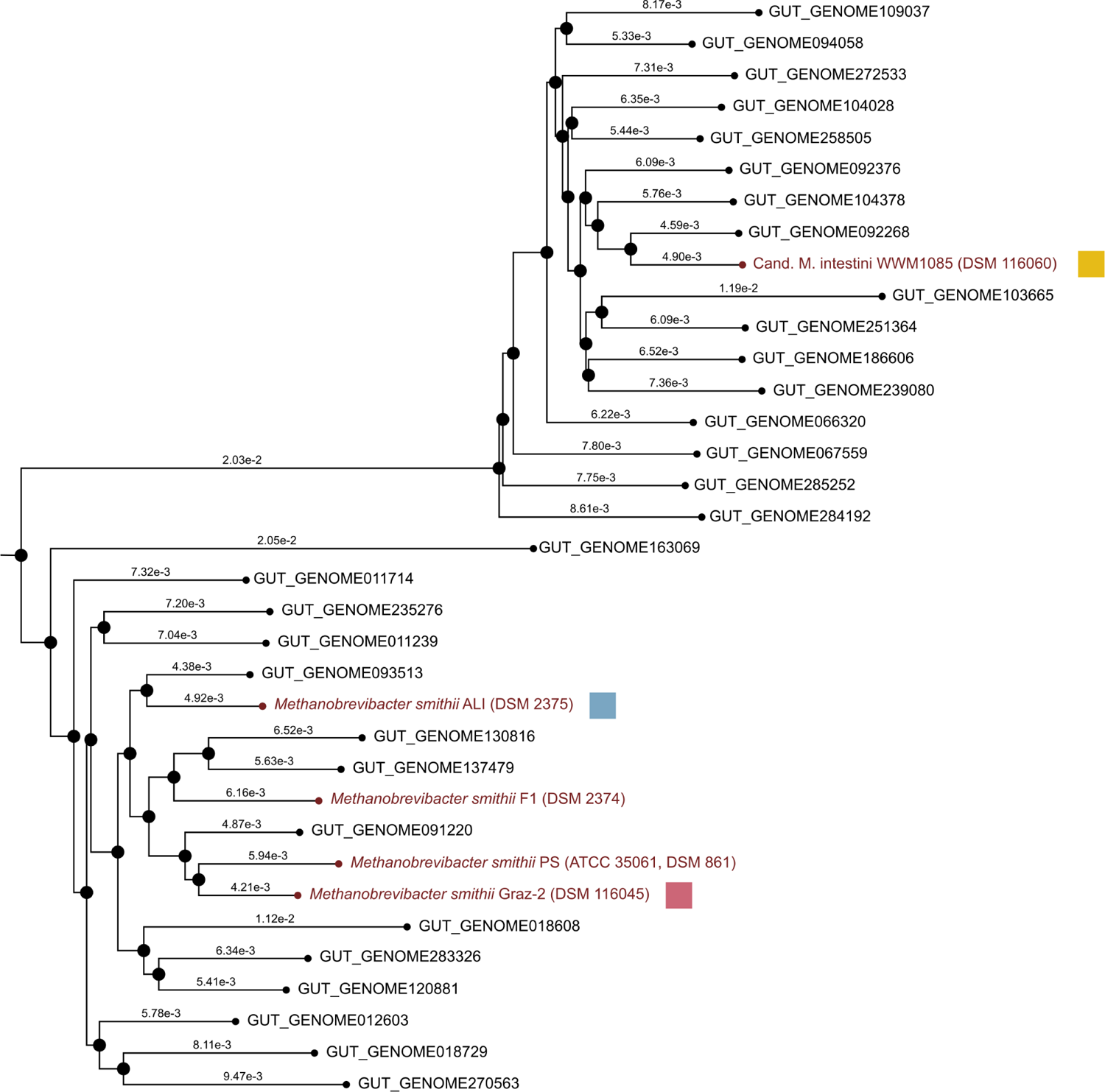
Neighbour Joining tree, calculated for genomes of *Methanobrevibacter* isolates (which are available in culture collections and are shown in dark red), and respective MAGS (Chibani et al., 2022) from the *M. smithii* clade. Representative genomes of the recently identified clade centered around WWM1085 (highlighed with a yellow square) are designated as “Mbb_smithii_A” based on the current GTDB classification. Consistent and stable clustering of the two *Methanobrevibacter* clades was observed. Pink square indicates *M. smithii* GRAZ-2, blue square *M. smithii* ALI. Distances based on the Mash distance matrix (24) are correlated to the average nucleotide identity (ANI) such as D ≈ 1-ANI.

WWM1085 shares its closest genetic relationship with the readily accessible *M. smithii* strain, known as *M. smithii* ALI. Notably, this particular strain was isolated from human gastrointestinal samples as well (in contrast to the available *M. smithii* PS, which was isolated from sewage). Consequently, throughout this study, WWM1085 and *M. smithii* GRAZ-2 were specifically compared to *M. smithii* ALI.

### Polar lipid composition and lipid-associated sugars

Major detected lipids were largely congruent across species and strains, with archaeol (C_43_H_88_O_3_) being the most prevalent lipid (Relative abundance for *M. smithii* ALI: 83.93%; WWM1085: 72.57%; and *M. smithii* GRAZ-2: 93.85%), followed by Caldarchaeol (C_86_H_172_O_6_) (*M. smithii* ALI: 13.65%; WWM1085: 26.37%; *M. smithii* GRAZ-2: 5.81%), and cyclic arachaeol (C_43_H_86_O_3_) (*M. smithii* ALI: 0.65%; WWM1085: 1.06%; *M. smithii* GRAZ-2: 0.34%). Traces of glycerol dialkyl glycerol tetraether lipids (GDGT-1) or H-shaped caldarchaeol were found in *M. smithii* ALI (1.77%), but not in the other strains. Lipid-associated sugar profiles were very similar for all strains with glucose being most prevalent, accompanied by minor amounts of fructose, rhamnose, ribose and xylose.

### Comparative genomics and metabolomics

Detailed genomic comparisons of WWM1085 with available *M. smithii* genomes are provided in our earlier publication (8), indicating several differences. For instance, WWM1085 does not possess modA/B for molybdate transport. The *M. smithii*_A genomes (including those from MAGs) were further characterized by additional unique membrane/cell-wall-associated proteins, such as adhesin-like proteins, surface proteins and a number of uncharacterized membrane proteins/transporters (8). Notably, *M. smithii* GRAZ-2 and the WWM1085 genome contained the ABC.FEV.P/S/A iron transport system (EC 3.6.3.34), which was distinctive to all other tested genomes, indicating a potential adaptation towards the human gut environment, where iron is a highly-demanded resource.

Utilizing NMR-based metabolomics, the turnover of metabolites was examined among three strains (*M. smithii* ALI, *M. smithii* GRAZ-2, and WWM1085) in a medium containing yeast extract. All strains reached the stationary phase after 72 h (*M. smithii* ALI and WWM1085) or at the latest, after 168 h (*M. smithii* GRAZ-2) of growth (growth curves shown in Supplementary Fig. 1).

All strains exhibited a notable and expected statistically significant uptake of acetate and production of succinate, which was highest in the WWM1085 culture (4-fold increase; Fig. 4) (Supplementary Table S7).

**Fig. 4.**
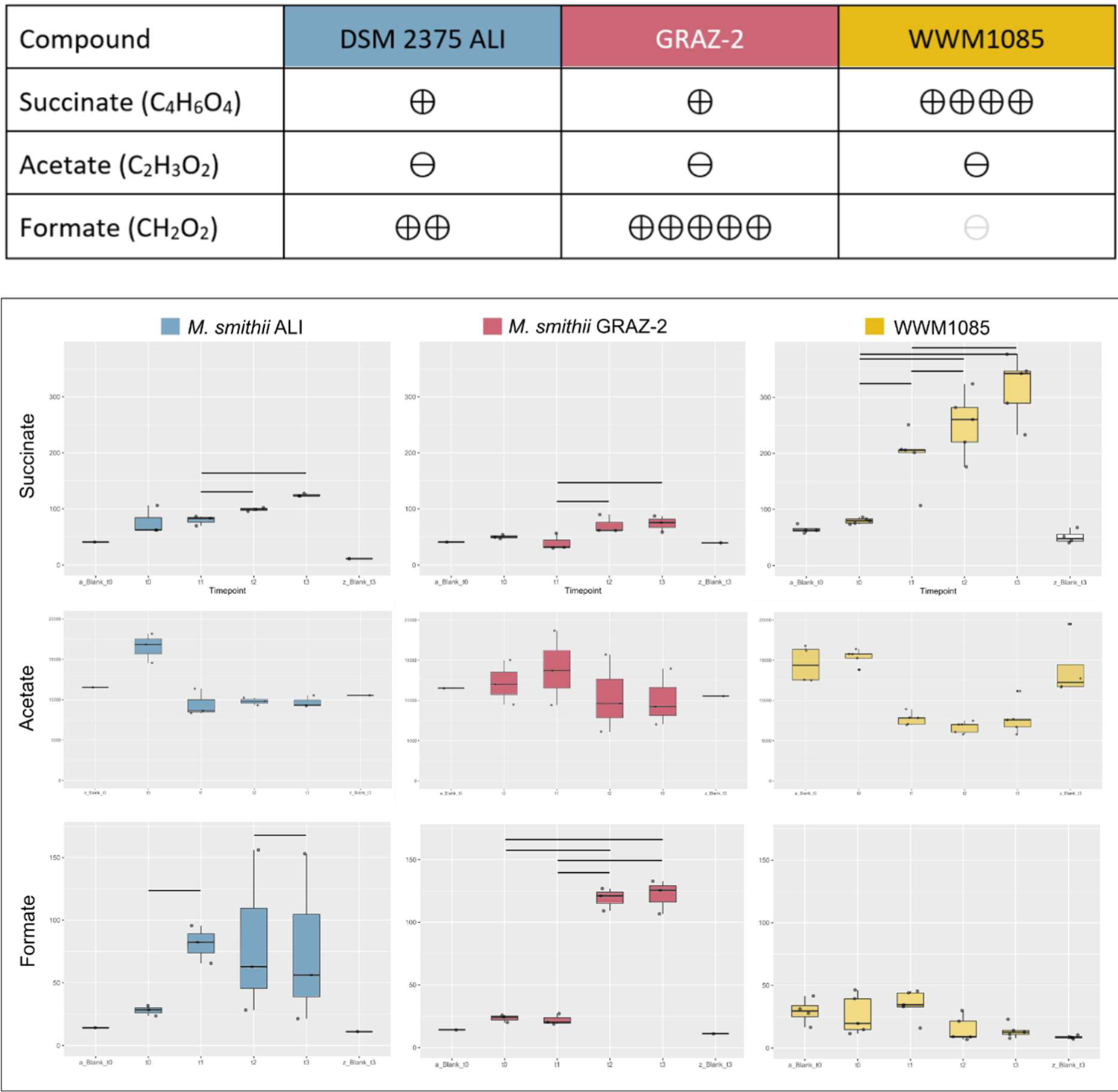
Metabolic dynamics and variability in various compounds among the studied *Methanobrevibacter* strains in MS medium supplemented with yeast extract. Concentrations in µM/L. Upper panel (table): The symbols indicate statistically significant changes over time, with the number of symbols reflecting the fold change (e.g., 5 symbols denote a 5-fold change or more). The gray symbol signifies a statistical trend (p=0.05X). The symbol “-” denotes no change. Lower panel: Boxplots of the respective measurements (all original data provided in Supplementary Table S7). It’s noteworthy that media blanks did not exhibit any statistically significant changes in compound levels.

Unlike the *M. smithii* cultures, WWM1085 did not exhibit formate accumulation in the medium. Specifically, there was a notable and substantial increase in formate accumulation for *M. smithii* GRAZ-2, reaching a fivefold increase (Fig. 4).

All biological properties of the tested strains, including *M. smithii* PS are provided in Table 2.

**Table 2.**
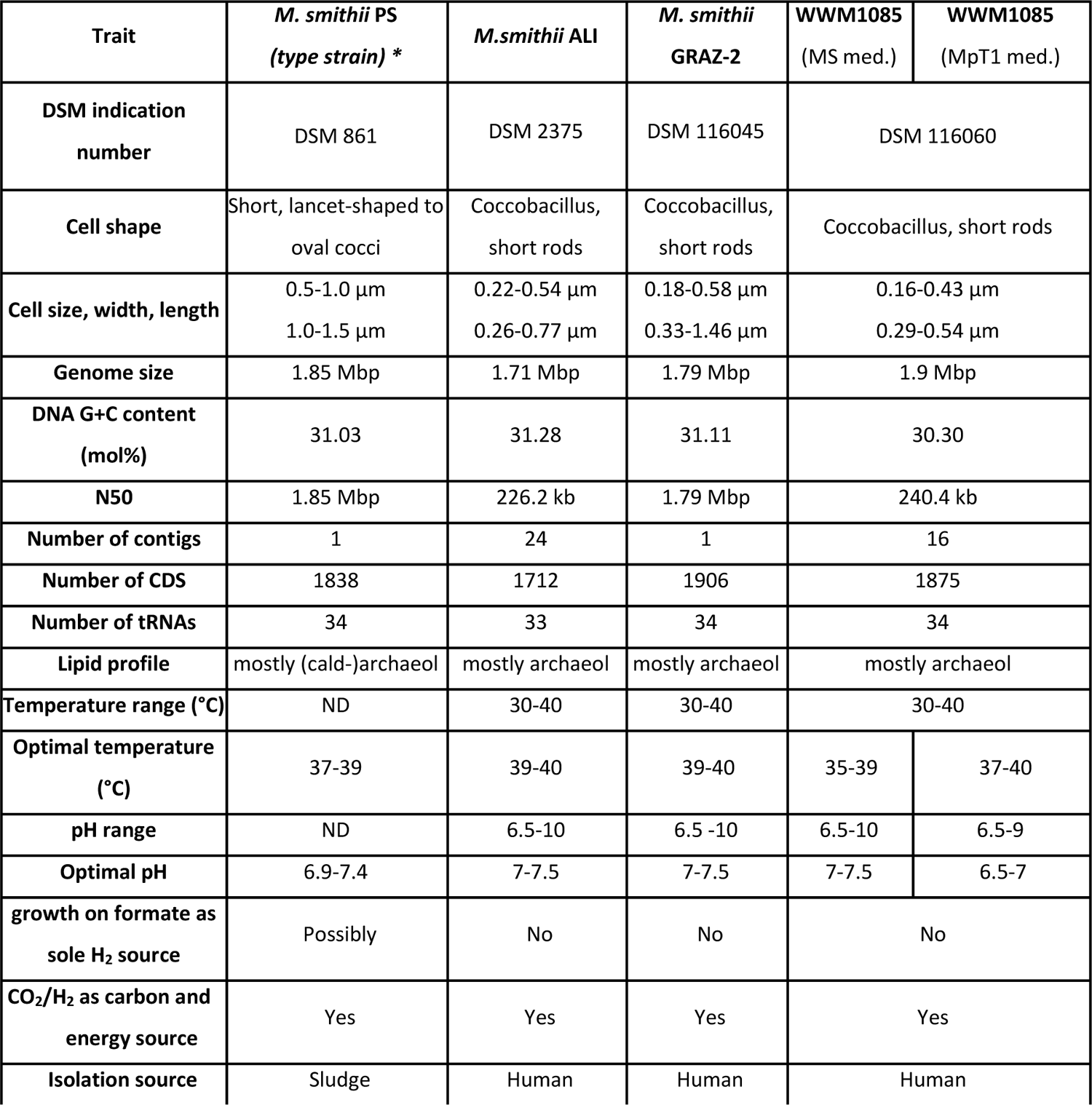
Distinguishing features among the strains *M. smithii* ALI, *M. smithii* GRAZ-2, and WWM1085. * Description for type strain *M. smithii* PS taken from (40) and (42). ND: Not determined, med: medium.

| Trait | *M. smithii* PS<br>(type strain) * | *M.smithii* ALI | *M. smithii*<br>GRAZ-2 | WWM1085<br>(MS med.) | WWM1085<br>(MpT1 med.) |
| --- | --- | --- | --- | --- | --- |
| DSM indication number | DSM 861 | DSM 2375 | DSM 116045 | DSM 116060 |  |
| Cell shape | Short, lancet-shaped to oval cocci | Coccobacillus, short rods | Coccobacillus, short rods | Coccobacillus, short rods |  |
| Cell size, width, length | 0.5-1.0 µm | 0.22-0.54 µm | 0.18-0.58 µm | 0.16-0.43 µm |  |
|  | 1.0-1.5 µm | 0.26-0.77 µm | 0.33-1.46 µm | 0.29-0.54 µm |  |
| Genome size | 1.85 Mbp | 1.71 Mbp | 1.79 Mbp | 1.9 Mbp |  |
| DNA G+C content (mol%) | 31.03 | 31.28 | 31.11 | 30.30 |  |
| N50 | 1.85 Mbp | 226.2 kb | 1.79 Mbp | 240.4 kb |  |
| Number of contigs | 1 | 24 | 1 | 16 |  |
| Number of CDS | 1838 | 1712 | 1906 | 1875 |  |
| Number of tRNAs | 34 | 33 | 34 | 34 |  |
| Lipid profile | mostly (cald-)archaeol | mostly archaeol | mostly archaeol | mostly archaeol |  |
| Temperature range (°C) | ND | 30-40 | 30-40 | 30-40 |  |
| Optimal temperature (°C) | 37-39 | 39-40 | 39-40 | 35-39 | 37-40 |
| pH range | ND | 6.5-10 | 6.5 -10 | 6.5-10 | 6.5-9 |
| Optimal pH | 6.9-7.4 | 7-7.5 | 7-7.5 | 7-7.5 | 6.5-7 |
| growth on formate as sole H<sub>2</sub> source | Possibly | No | No | No |  |
| CO<sub>2</sub>/H<sub>2</sub> as carbon and energy source | Yes | Yes | Yes | Yes |  |
| Isolation source | Sludge | Human | Human | Human |  |

### Description of Methanobrevibacter intestini sp. nov

*Methanobrevibacter intestini* (in.tes.ti’ni. L. gen. n. *intestini*, of the gut). Coccobacillus with slightly tapered or rounded ends, about 0.16-0.43 µm in width and 0.29-0.54 µm in length, occurring mostly in pairs or short chains. The DNA GC content is 30.30 mol%. Optimum temperature: 35-40°C; optimum pH 6.5-7.5. Strictly anaerobic. Grows and produces methane from H_2_ and CO_2_. Requires acetate and additional organics (e.g. yeast extract) for growth. Can not grow on formate as a sole electron source. Genome comparisons with type species *M. smithii* PS revealed numerous differences including an average nucleotide identity of 93.55%. With such, WWM1085 represents a novel species within the *Methanobrevibacter* genus, and is the first isolated representative. Strain WWM1085 was isolated from human feces from an US American individual. The type strain WWM1085^T^ (=DSM 116060, CECT 30992). The GenBank accession number of its genome is NQLD00000000.

### Sequencing data

The GenBank accession number of the genome of WWM1085 is NQLD00000000. The genome of GRAZ-2 is available through BioProject ID PRJNA1067514. The 16S rRNA gene sequence of WWM1085 is available through NCBI GenBank PP338268.

## Acknowledgements

We would like to acknowledge the computational resources of the MedBioNode at the Medical University of Graz, as funded by the Austrian Federal Ministry of Education, Science and Research, Hochschulraum-Strukturmittel 2016 grant as part of BioTechMed Graz, and the support of the ZMF team at the Core Facility Computational Bioanalytics (Medical University of Graz). We further thank Birgit Grün and Gesa Martens (DSMZ) for excellent technical assistance.

