## Supplementary Information for "Expanding the cultivable human archaeome: *Methanobrevibacter intestini* sp. nov. and strain *Methanobrevibacter smithii “*GRAZ-2*”* from human feces"

Viktoria Weinberger[1†], Rokhsareh Mohammadzadeh[1†], Marcus Blohs[1], Kerstin Kalt[1], Alexander Mahnert[1], Sarah Moser[1], Marina Cecovini[1], Polona Mertelj[1], Tamara Zurabishvili[1], Jacqueline Wolf[3], Tejus Shinde[1], Tobias Madl[2,4], Hansjörg Habisch[4], Dagmar Kolb[5,6], Dominique Pernitsch[5], Kerstin Hingerl[5], William Metcalf[7], Christine Moissl-Eichinger[1,2*]

[1] D&R Institute of Hygiene, Microbiology and Environmental Medicine, Medical University of Graz, Graz, Austria
[2] BioTechMed Graz, Graz, Austria
[3] Research Group Metabolomics, Leibniz Institute DSMZ-German Collection of Microorganisms and Cell Cultures GmbH, Braunschweig, Germany.
[4]Otto Loewi Research Center, Medicinal Chemistry, Research Unit Integrative Structural Biology, Medical University of Graz, Graz, Austria
[5] Core Facility Ultrastructure Analysis, Medical University of Graz, Graz, Austria
[6] Gottfried Schatz Research Center, Cell Biology, Histology and Embryology, Medical University of Graz, Graz, Austria
[7]Department of Microbiology, University of Illinois, Urbana, Illinois, USA

*Corresponding author


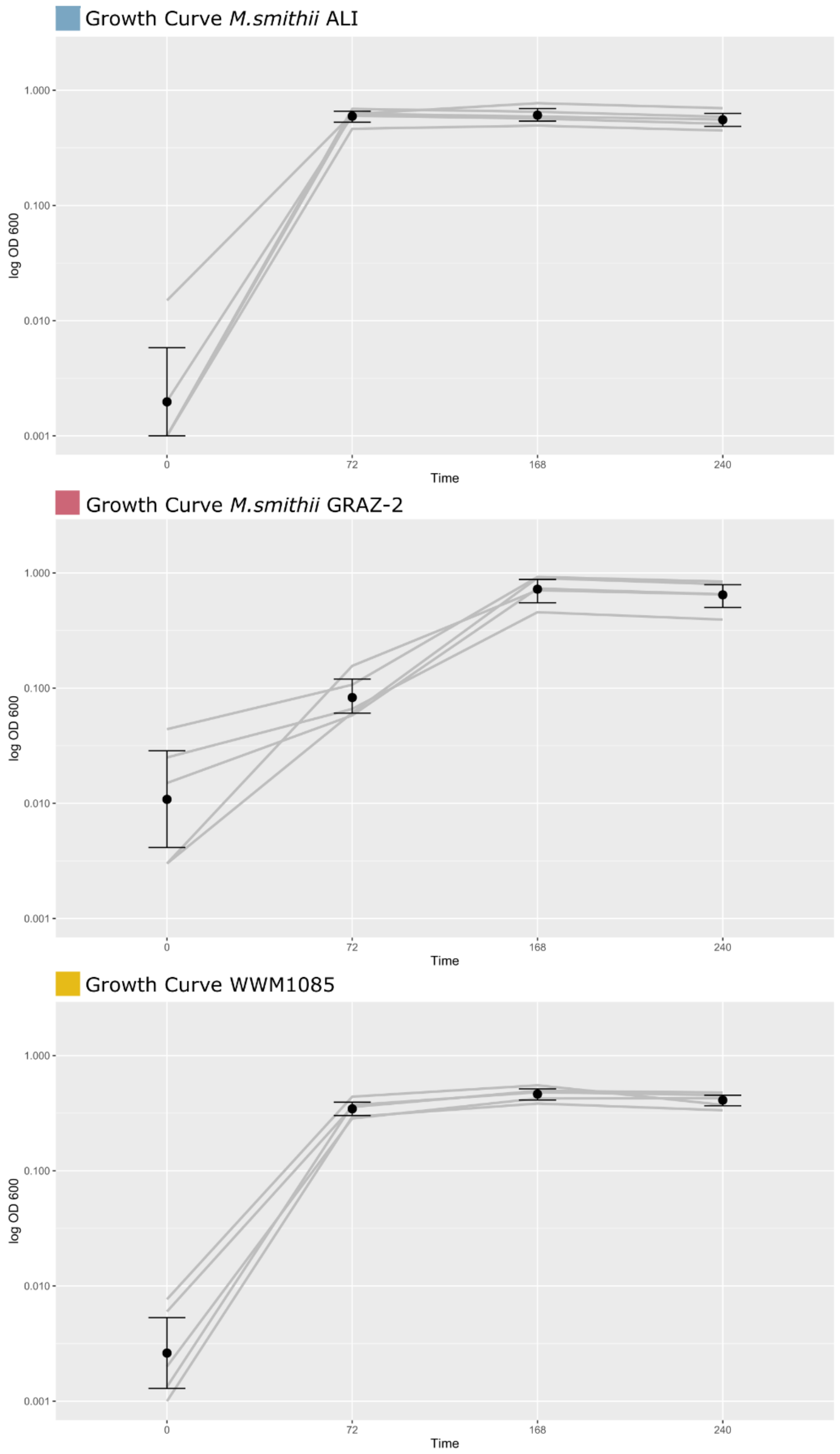


**Supplementary Fig. 1**: Growth curves of all strains. X-axis indicates time in hours, y-axis displays the log $OD_{600}$. *M. smithii* ALI and WWM1085 reached the stationary phase at 72 h, while *M. smithii* GRAZ-2 reached the stationary phase after 168 h.

**All Supplementary Tables are provided in the following Github repository**:

https://github.com/Christine-Moissl-Eichinger/Methanobrevibacter_description

This repository contains:

**Supplementary Table S1: Full length 16S rRNA gene sequences of Methanobrevibacter isolates.**

**Supplementary Table S2: Distance Matrix of 16S rRNA genes**

**Supplementary Table S3: mcrA genes of Methanobrevibacter isolates.**

**Supplementary Table S4: Distance Matrix of mcrA genes**

**Supplementary Table S5: Genomes and all associated information used herein for comparisons.**

**Supplementary Table S6: ANI values of full genome comparisons.**

**Supplementary Table S7: Metabolomics, concentrations in uM/L**
